# Early language dissociation in bilingual minds: Non-invasive magnetoencephalography evidence through a machine learning approach

**DOI:** 10.1101/2023.03.22.533825

**Authors:** Nicola Molinaro, Sanjeev Nara, Manuel Carreiras

**Affiliations:** Basque center on Cognition, Brain and Language, Donostia/San Sebastian, Spain; Ikerbasque, Basque Foundation for Science, Bilbao, Spain; Mathematical Institute, Department of Mathematics and Computer Science, Physics, Geography, Justus-Liebig-Universität Gießen (University of Giessen), Gießen, Germany; University of the Basque Country. UPV/EHU

**Author notes:** Equal contribution as first author (Nicola Molinaro, Sanjeev Nara).

**Keywords:** Bilingualism, Neural decoding, Magnetoencephalography, Lexical processing, Word Reading, Picture Naming

## Abstract

Is language selection in balanced bilinguals decodable from neural activity? Previous research employing various neuroimaging methods has not yielded a conclusive answer to this issue. However, direct brain stimulation studies in bilinguals have detected different brain regions related to language production in separate languages. In the present MEG study, we addressed this question in a group of proficient Spanish-Basque bilinguals (N=45), who performed two tasks (picture naming and word reading). They were asked to name the line drawing or read the word out loud, either in Basque or Spanish, if the ink turned from black to green after one second (randomly, in 10% of trials). Sensor-level evoked activity was similar and could not be differentiated for the two languages in either task. Crucially however, decoding analyses classified the language used in both tasks, starting ∼100 ms after stimulus onset. Searchlight analyses revealed that activity detected in the right occipital-temporal sensors contributed the most to language decoding in the picture naming task, while the left occipital-temporal sensors contributed the most to decoding in word reading. The cross-task decoding analysis highlighted robust generalization effects from the picture naming to the word reading task in a later time interval. The present findings bridge the gap between non-invasive and invasive experimental evidence on bilingual lexical processing and provide novel evidence about the role of the two hemispheres in activating each language for picture naming and word reading.

## Introduction

Bilingual language processing implies dealing with different linguistic codes at the same time: the bilingual speaker must keep both languages ready for efficient use in real life, regardless of whether information from one language is segregated from the other or whether both languages are represented together. This astonishing ability has continually attracted neuroscientists to the challenge of determining if distinct neural substrates are recruited by the use of different languages in the brain (Fabbro, 2001; Ojemann and Whitaker, 1978). However, evidence from non-invasive neuroimaging has not provided a conclusive answer to this issue.

A number of recent meta-analyses have explored whether there is supporting evidence for distinct neural correlates of the processing of different languages in the bilingual brain (for a recent overview, see Comstock and Oliver, 2021). Overall, the reviewed evidence highlights that the core cortical network activated by words in different languages is the same, with relatively minor differences that could be explained by differences in proficiency in the two languages, the use of those languages, or the relative age of acquisition. When considering balanced bilinguals, highly proficient in both languages, no differences are typically observed across various neuroimaging techniques: left hemisphere frontotemporal regions are activated across languages and tasks (Hernandez et al., 2000; Jones et al., 2012), as well as, to a lesser extent, homologous regions of the right hemisphere (e.g., Geng et al., 2022; Nichols et al., 2021), especially in lexical/semantic tasks (Sulpizio et al., 2020). This “null effect” supports the idea that the neural infrastructure supporting the two languages is not differentiated and, consequently, that the lexical repertoire of the bilingual mind is integrated and indistinct for the two languages. This “integrated view” is supported by evidence that the use of words in one language mentally activates the words of the other language even in cases where the two linguistic codes are highly different (Emmorey et al., 2008; Morford et al., 2011; Thierry and Yan, 2007; Villameriel et al., 2022).

In addition, ERP studies highlighted an early influence on target language word processing from lexical neighbors of the other language (e.g., Midgley et al., 2011; Peeters et al., 2013). This indicates that, during reading, language selection is a relatively late process, preceded by earlier lexical activation dynamics that are not language specific. Along these lines, the Bilingual Interactive Activation + (BIA+) model (Dijkstra and van Heuven, 2002) assumes a high level of cross-language interaction in lexical processing by balanced bilinguals. Bilingual lexical processing does not seem to recruit highly segregated mental representations or tap into separate cognitive mechanisms for the two languages. These points support the observation that the neural underpinnings of bilingual language processing do not necessarily need to rely on functionally and spatially separated networks.

However, evidence against the “integrated view” has been provided by direct brain stimulation research (Lucas et al., 2004; Ojemann and Whitaker, 1978). During awake brain stimulation, it is typically observed that stimulation of different areas can selectively impair the use of one language or the other. Based on this evidence, the idea that different functional brain networks are recruited for separate languages should be reconsidered. One possibility is that the same macro-level brain systems are recruited for both languages, while differences could emerge at a more micro-level: highly overlapping functions would be recruited for the neural computation of the two languages, yet language segregation could emerge by better scrutinizing the trial-by-trial variability in the neural correlates. Classical univariate analysis would not be suited for exploring such variability since the individual brain responses are averaged across multiple trials and the fine-grained trial-by-trial variability is lost. A similar rationale has been put forward by Xu and colleagues (2017), who explored the multivariate pattern of fMRI responses to different languages in a group of bilinguals. Instead of performing univariate analysis they employed a multivariate approach and observed functional separation for the two languages across regions of the language brain network. This methodological approach inspired the present investigation, in which we focused on the functional dissociation between the time-courses of the neural responses to the two languages of balanced bilinguals.

In the present MEG (magnetoencephalography) study, we used a picture naming and a word reading task to assess covert production of lexical information within the same group of Basque-Spanish balanced bilinguals. We employed a classic evoked activity analysis approach (*ERFs, Event-Related Fields*) to compare the electrophysiological correlates of picture naming and word reading in the two languages. More importantly, we used a machine learning approach (Cichy et al., 2014; Grootswagers et al., 2017; King and Dehaene, 2014; Nara et al., 2021a) to explore the trial-by-trial variability in the two languages and evaluate a possible more fine-grained functional language classification in the neural responses. This analysis reveals, across time-points, if the neuromagnetic correlates of the two languages can be accurately decoded, thus showing when the neural activity becomes language specific (*temporal decoding*). We then performed a searchlight analysis across the whole-time window to investigate the sensor locations contributing maximally to the decoding. We also analyzed if the decoder trained at a specific time-point could generalize across the other time points (*temporal generalization*). This analysis can reveal if separate sources of variability explain decoding accuracy at two separate time points - evidence of separate neurocognitive processes - or if the same source of variability underlies decoding accuracy at both time points - evidence of the same neurocognitive processes. Finally, we explored if the decoder trained at a specific time point for a specific task (either picture naming or word reading) could generalize across the time points of the other task (*cross-decoding*). This approach will reveal if language-specific representations at a specific time in one task could explain brain activity across time points in the other task. Overall, we expect the decoding approach to provide more precise information concerning bilingual lexical processing compared to classical evoked activity analyses. Based on the integrated view, no reliable decoding should be observed in any task. In contrast, if the two languages recruit different neural resources, we should observe significant language classification. Previous decoding studies (Dirani and Pylkkänen, 2023; Giari et al., 2020; Leonardelli et al., 2019) reported earlier decoding effects in language tasks for pictures (starting ∼150 ms), than for words (after 200 ms). Given this evidence that semantically meaningful stimuli can modulate neural activity so early, we can expect language decoding to emerge in these early time windows as evidence of early language selection processes.

## Methods

### Participants

Forty-five Spanish-Basque bilinguals (29 females; age range: 19-42; mean age: 27.1 years; standard deviation: 5.2) took part in the present study. All of them were right-handed and had normal or corrected-to-normal vision. All participants acquired both languages (Spanish and Basque) before three years old. All participants went through a short semi-structured oral proficiency interview in each language (de Bruin et al., 2017) on which they all scored higher than 4 (on a 5-point scale) in both languages. In a picture naming test, across a set of 65 items per language, participants correctly named 64.6 pictures in Spanish (sd: 0.83; range 61-65) and 61.8 pictures in Basque (sd: 2.86; range 56-65), showing a high level of proficiency in both languages. All participants gave their written informed consent in accordance with ethical guidelines approved by the Research Committees of the Basque Center on Cognition, Brain and Language (BCBL).

### Stimuli

We used a set of 321 line-drawn pictures representing concrete nouns that have been used in previous studies (e.g., Rueckl et al., 2015). From this initial set, we selected 288 pictures (mainly determined by the exclusion of pictures referring to phonologically similar words in the two languages, i.e., cognates) that unambiguously referred to a specific noun (name agreement > 80%, based on Gisbert-Muñoz et al., 2021).

Reference databases (Spanish: EsPal, Duchon et al., 2013; Basque: E-Hitz, Perea et al., 2006) largely differed in corpus size, however these resources still provide useful information about the words referring to each picture in both languages. When considering all 288 items, despite using translation equivalents for each picture, there were slight significant differences between the languages on a number of relevant variables. Spanish words had slightly higher frequency values (LogFreq: mean=1.0, sd=0.6) than Basque words (LogFreq: mean=0.9, sd=0.7), and were shorter (number of letters: mean=5.9, sd=1.8) than Basque words (mean=6.3, sd=2). In addition, words in Spanish had fewer syllables (mean=2.5, sd=0.8) compared to Basque (mean=2.7, sd=0.9). It is important to remark that we used the same words in the two languages. While the frequency parameters are not completely comparable for the two datasets (given the difference in the size of the reference corpora), the length effects were expected since Basque has a richer agglutinating morphology than Spanish. We further considered this issue by conducting, where possible, complementary analyses with a subset of strictly balanced items. Specifically, we split our stimuli into two sets of high and low frequency items in the two languages (see Table 1 and Supplementary Figure 3). Of note, high frequency items (but not low frequency ones) were closely balanced across languages for the three parameters of interest.

**Table 1:** Lexical parameters of the critical words in the two languages separated by lexical frequency.

|  | Spanish | Basque | t-value |
| --- | --- | --- | --- |
| **High Frequency items** |  |  |  |
| Log frequency | 1.52 (0.43) | 1.49 (0.40) | 0.53 |
| Number of letters | 5.41 (1.56) | 5.38 (1.59) | 0.19 |
| Number of syllables | 2.29 (0.64) | 2.32 (0.69) | 0.79 |
| **Low Frequency items** |  |  |  |
| Log frequency | 0.54 (0.27) | 0.34 (0.32) | 5.82*** |
| Number of letters | 6.33 (1.84) | 7.24 (2.04) | 4.01*** |
| Number of syllables | 2.64 (0.79) | 3.08 (0.89) | 3.91** |

*\* $p < 0.01$ , \*\* $p < 0.001$ , \*\*\* $p < 0.0001$ , extracted from a Welch Two Sample $t$ -test*

These items (a picture and the corresponding words in each language) were used to create six sets of 96 stimuli for which we balanced the overall lexical properties (in both Spanish and Basque) of the target nouns referring to each picture. Across sets, each item was present twice. From these six sets, four of them were assigned to each task and condition (naming or reading, either in Spanish or in Basque) determining six combinations (Latin square design) so that across participants each picture was named in each language and each word was read in each language the same number of times.

### Procedure

Stimuli were delivered using Psychtoolbox 3 (Brainard, 1997) in Matlab 2014b. Visual stimuli were presented in a MEG room on a back-projection screen with an external projector at 60 Hz. The screen was placed at 60 cm from the participant’s head, so that the stimuli covered a maximum visual angle of 5 degrees at the center of the screen. Participants performed the picture naming and word reading tasks separately in one session. Each task was divided into 16 blocks of 12 trials in one of the two languages. To ensure participants understood the instructions, they performed 20 practice trials under the experimenter’s observation at the start of each task (10 in each language). Overall, including participant preparation, the whole experimental session lasted ∼2 hours.

In the picture naming task, a fixation point was presented at the center of the screen for a jittered interval of 2500 ms on average (± 1500 ms) followed by a picture drawn in black ink for an additional 1000 ms. On 10% of trials the ink turned to green, prompting the participant to name the picture out loud. Otherwise, the fixation point reappeared on the screen for the next trial. Trials requiring overt speech production were not considered in the analysis. The covert naming procedure was employed to avoid pre-articulatory and motor artifacts contaminating MEG recordings in a potentially theoretically critical time interval. Naming latencies could reach 350 ms, thus affecting the time intervals typically associated with lexical-semantic processing (Kutas and Federmeier, 2011).

Each block of twelve trials was preceded by a Spanish or Basque flag to indicate the language that should be used for naming. At the end of each block a self-paced break was included. Naming accuracy was monitored by the experimenter in the control room (each participant’s accuracy > 91%).

In the word reading task, we used the same experimental structure (jittered fixation point followed by the stimulus at the center of the screen). However, instead of pictures, words printed in black (turning to green in 10 % of cases) were presented (font: Lucida Console; size: 34). The same instruction at the beginning of each block informed the participant about the language of the next trials. Reading accuracy was monitored by the experimenter in the control room (each participant’s accuracy > 94%).

### Magnetoencephalographic recordings

MEG data were acquired in a magnetically shielded room using the whole head MEG system (MEGIN-Elekta Neuromag, Finland) installed at BCBL (https://www.bcbl.eu/en/infrastructure-equipment/meg). The MEG system consists of 306 sensors (102 magnetometers and 204 gradiometers arranged in a helmet configuration). The head position during the experiment was continuously monitored with five Head Positioning Indicator coils (HPI). The location of HPI coils were defined relative to three fiducial points (nasion, left preauricular (LPA) and right preauricular (RPA) point) using 3D head digitizer (Fastrack Polhemus, Colchester, VA, USA). This process is critical for correction of movement artifacts (movement compensation) during data acquisition. The MEG data were recorded continuously with a bandpass filter (0.01 – 330 Hz) and a sampling rate of 1 kHz. Eye movements were monitored with two pairs of electrodes placed above and below each eye (horizontal electrooculogram - HEOG), and on the external canthus of both eyes (vertical electrooculogram - VEOG). Similarly, electrocardiogram (EKG) was also recorded using two electrodes (bipolar montage) placed on the right side of the abdomen and below the left clavicle of the participant. The continuous MEG data were pre-processed offline to suppress the external electromagnetic field using temporal Signal-Space-Separation (tSSS; Taulu and Simola, 2006) method implemented in Maxfilter software (MaxFilter 2.0). The MEG data were also corrected for movement compensation and bad channels were repaired within the MaxFilter algorithms. All the analyses reported below were carried out in Matlab version 2021B.

## Data analysis

### Event-Related Fields

Continuous MEG data were first segmented into time intervals starting 1 second before the onset of each visual stimulus until 2 seconds afterwards. Based on visual inspection, we detected epochs containing large artifacts (SQUID jumps, large head movements) and excluded them from the following analyses. On average 3% of trials were rejected, with no differences among conditions. A FastICA algorithm (Hyvärinen, 1999) was then used to detect independent components which can represent either eye-movements or heart-beats. Across participants we excluded 1 to 3 components based on visual inspection. We focused on planar gradiometers for the evoked analysis. After baseline correction (−500 to −10 ms), gradiometer pairs in the same location were combined using the root-mean square approach. Afterwards, we separated the trials based on conditions for averaging, independently for each participant. Potential peaks of interest were select as the maximum value in a time interval that exponentially increased across time, ranging from 20 ms to 500 ms.

Statistical analyses were performed comparing the neural activity in the two languages (Spanish and Basque) independently for each task (picture naming and word reading) using the cluster-based approach (10000 permutations, cluster alpha: p<0.05, alpha: p<0.05) on the time interval of interest (0-1000 ms) across the whole set of sensors (Maris and Oostenveld, 2007).

### Time-resolved decoding

Time-resolved decoding (also called temporal decoding) was used to decode the language used by participants (Spanish or Basque) from neural patterns recorded by MEG. The preprocessed MEG data in both tasks (picture naming and word reading) were segmented between 200 ms prior to and 1000 ms after the onset of the stimuli. The data were first low pass filtered to 25 Hz and then down sampled to 200 Hz. A covariance-based trial rejection was performed to remove outlier trials. Trials deviating by less than 2 percent or more than 95 percent from the mean covariance were excluded prior to the classification procedure (average trial rejection ratio 7%). The data were then fed to the classifier separately for each task to decode the language of use, using a logistic regression classifier implemented in the MVPA Light toolbox (Treder, 2020). The Classification was performed separately at each time point for both tasks (words and pictures). We used k-fold cross-validation (k=5) to avoid overfitting and used the Area under the receiver operating characteristic (ROC) curve (AUC) for evaluating the classification performance. We also used an ‘*undersampling*’ approach to balance the number of items per condition before feeding the data for classification. This process was repeated 10 times for each subject to yield stable decoding patterns. The statistical significance of decoding across time was established using cluster corrected sign permutation tests (one-tailed) (similar to Dima et al., 2018; Nara et al., 2023, 2021b) applied to the AUC values obtained from the classifier with cluster-defining threshold (i.e., alpha: p<0.05; cluster-alpha: p<0.05; 10000 permutations).

### Searchlight across sensor-space

A searchlight analysis was implemented to determine the sensors contributing most to decoding the language from sensor-space MEG data. The data preprocessing steps (i.e., low pass filtering at 25 Hz, down-sampling at 200 Hz, logistic regression classifier, 5-fold cross validation, AUC as accuracy metrics) were the same as explained in the previous section. Unlike time-resolved decoding, the logistic regression classifier considered the whole-time interval (i.e., −200 ms to 1000 ms after the onset of stimulus presentation) to decode the language in both tasks separately (word reading and picture naming). To yield stable decoding patterns, the classification procedure was repeated 10 times for each subject. Since a longer time window was used, a more stringent threshold was applied to establish the statistical significance compared to time-resolved decoding. Cluster corrected sign permutation tests (one-tailed) were applied with cluster defining thresholds (i.e., alpha: p<0.001; cluster-alpha: p<0.001; 20000 permutations).

### Temporal Generalization

Time-resolved decoding helps to understand how cognitive processes unfold over time; however, this method does not reveal anything about the temporal organization of information processing stages in the human brain. The temporal generalization method measures the ability of a classifier to generalize across time (King et al., 2016; King and Dehaene, 2014). This method provides a novel way to understand how mental representations are manipulated and transformed for a given cognitive process. For this analysis, the data preprocessing steps (i.e., low pass filtering at 25 Hz, down-sampling at 200 Hz, logistic regression classifier, 5-fold cross validation) were the same as explained in the previous sections. Here a classifier is trained on each particular time point in the data and then tested over all other available time points separately for both tasks (i.e., picture naming and word reading). This results in a square matrix per block per participant where each point reflects the generalization strength of the classifier at a particular time (x axis) given the training at a given time point (y axis). To evaluate the presence of a group level effect, a cluster-based permutation test was used with cluster-defining threshold (i.e., alpha: p<0.05; cluster-alpha: p<0.05; 10000 permutations).

### Cross Decoding

Cross decoding is a variant of temporal generalization which generalizes across different experiments, tasks, or blocks. This method is used to understand whether the brain activity elicited in one cognitive scenario can predict brain activity in another scenario. In this study, the classifiers were first trained using the data from the picture naming task and then tested over the time points from the word reading task; this process was then repeated in reverse (classifiers trained on reading data and then tested on naming data). The parameters for classification were the same as explained in the previous three sections.

## Results

### Evoked activity

In the picture naming task, ERFs (shown in Figure 2), computed separately for the two languages, showed peaks of activity at around 130 ms in the occipital sensors, around 250 ms in occipito-temporal sensors, bilaterally, and around 450 ms in occipital and temporal sensors. Cluster-based permutations (Spanish vs. Basque) did not reveal any significant differences between the two conditions. However, visual inspection revealed that the early peak around 130 ms appeared to be reduced for Spanish compared to Basque. We therefore ran a post-hoc analysis in a time interval between 100 and 150 ms. The cluster-based approach indicated a negative cluster but this was not statistically significant (p=0.23).

**Figure 1:**
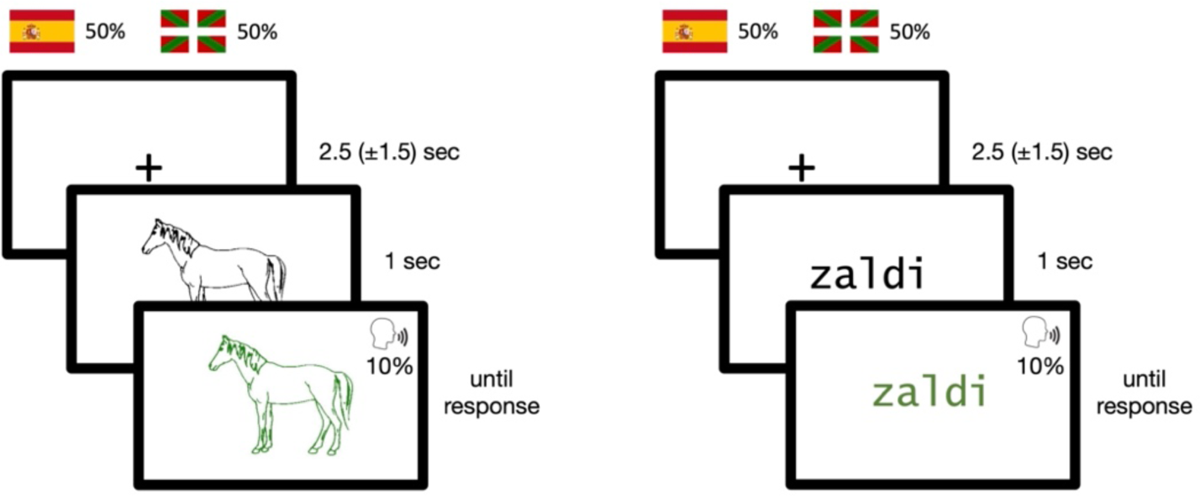
Example of one trial for the picture naming task (left) and the word reading task (right).

**Figure 2:**
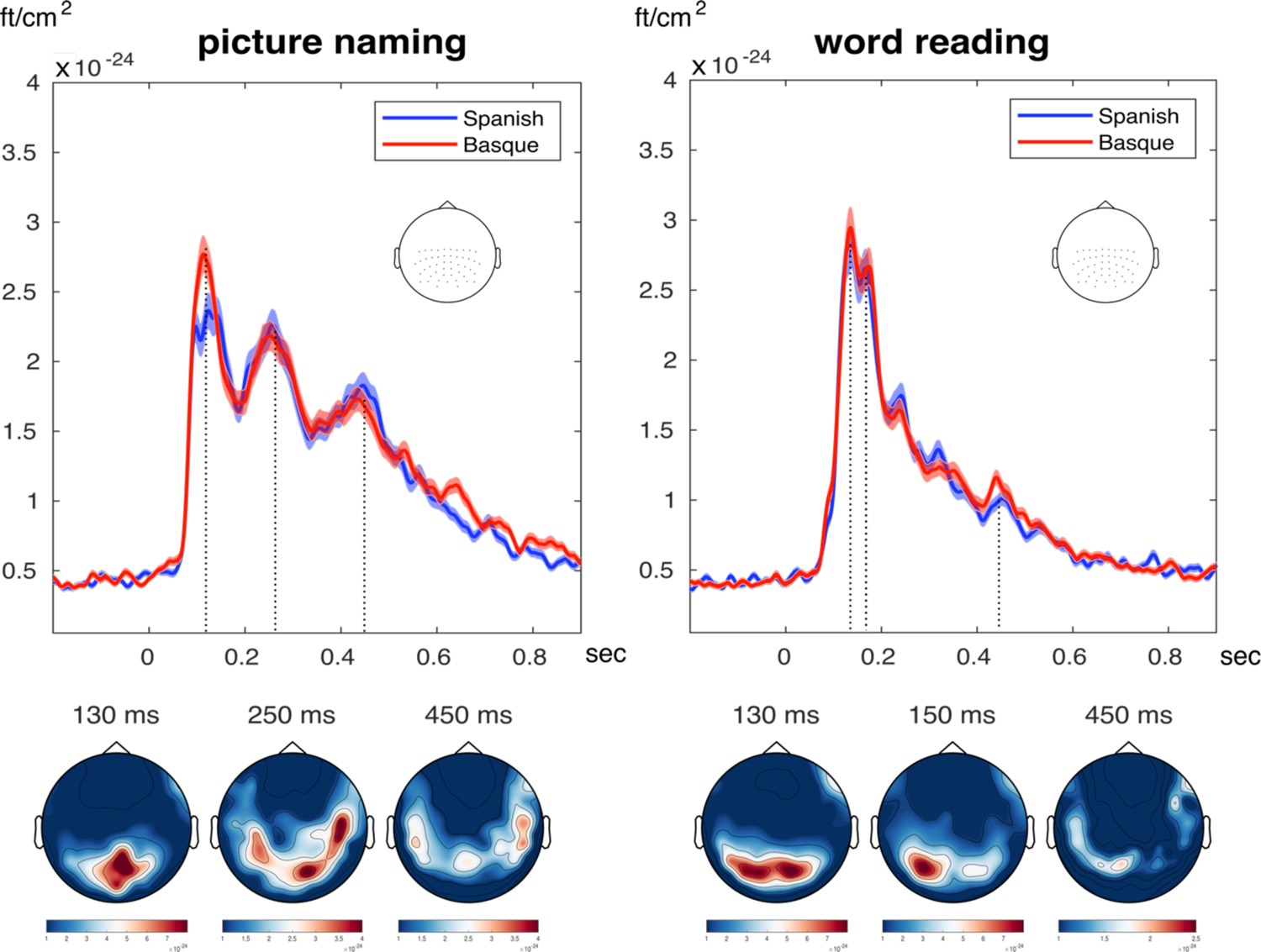
The top panels show event-related fields (combined gradiometers representation, averaged across a set of posterior sensors) elicited by Spanish (blue) and Basque (red) in the two experimental tasks. Shaded bars reflect the standard error from the mean. The bottom panels show topographies reflecting the sensors with maximum activity at representative time points marked with dotted lines in the time-course above.

In the word reading task, two early peaks could be clearly observed: at around 130 ms in the occipital sensors and at around 150 ms in the posterior left occipito-temporal sensors. A later peak was observed at around 400 ms, mainly evident in the left temporal sensors. Cluster-based permutations (Spanish vs. Basque) did not reveal any significant differences between the two conditions. Visual inspection showed a potential difference starting around 400 ms. A post-hoc analysis in a time interval between 400 and 500 ms indicated a negative cluster but this was not statistically significant (p=0.37).

Overall, the ERF data indicate that similar temporal dynamics emerged for Spanish and Basque across the whole stimulus set and for both tasks. This suggests that processing in the two languages is not characterized by any temporal differences.

### Time-resolved decoding

In contrast with the ERF data, neural representations of language (Spanish or Basque) were successfully decoded at above chance levels in both picture naming and word reading tasks. We observed one significant cluster (p<0.001) for both blocks (Figure 3). The language specific representations dissociated slightly earlier in picture naming (starting at 82 ms) compared to word reading (starting at 117 ms) and were sustained until the end of the time window considered. We also observed differences in peak decoding in the two tasks. The decoding for word reading peaked at 162 ms after the presentation of the stimulus whereas the peak decoding for the picture naming was at 432 ms. Thus, the decoding in the word reading task emerged slightly later but reached the peak earlier whereas in picture naming the decoding emerged earlier and slowly increased until ∼300-400 ms.

**Figure 3:**
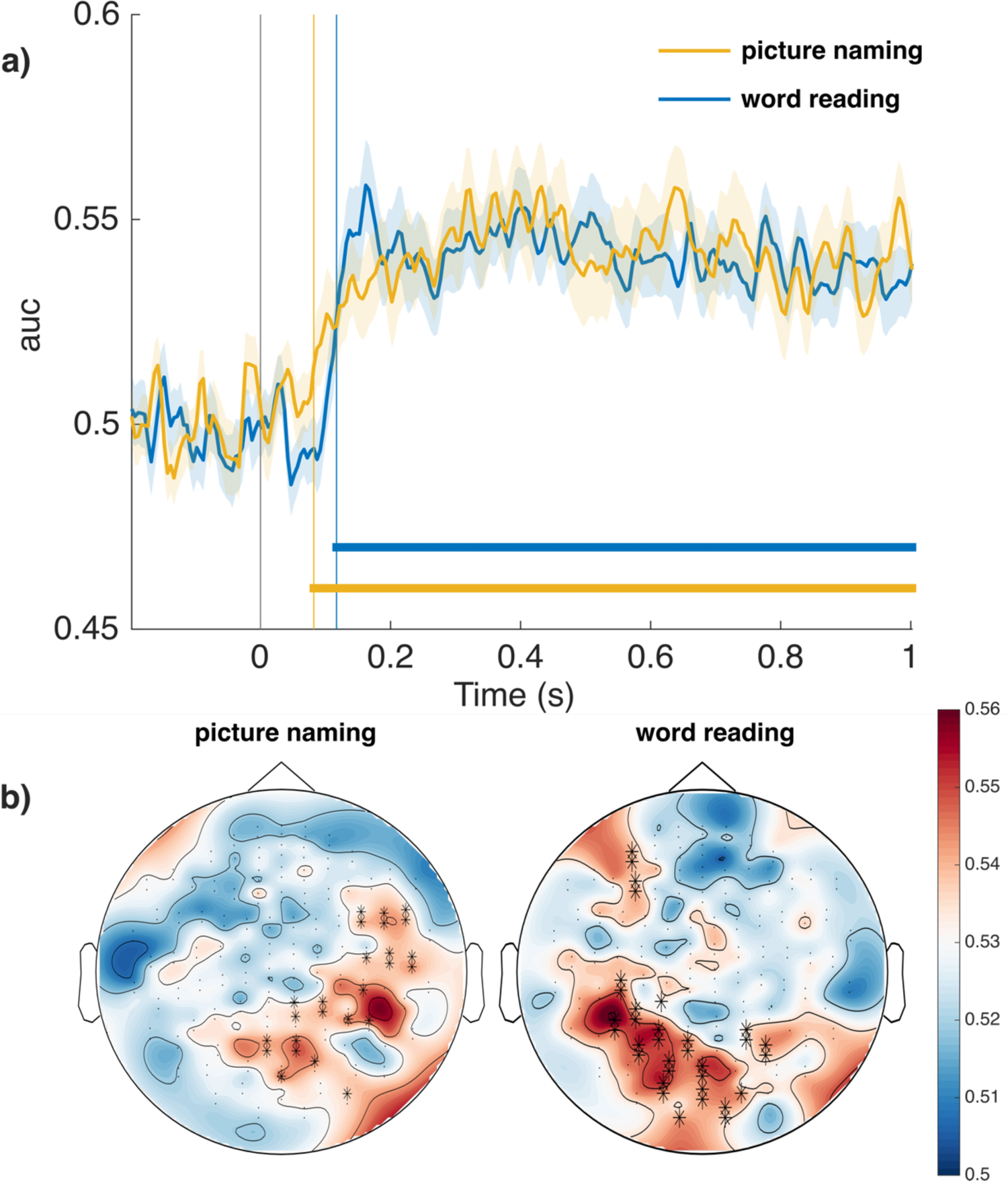
a) Time-resolved language decoding (Spanish vs. Basque, x-axis reports time in seconds) in word reading (blue) and picture naming (yellow). The horizontal lines in the figure indicate time points showing significant decoding above chance level (p < 0.05). b) Searchlight analysis of language decoding (Spanish vs. Basque) in word reading (left) and picture naming (right). The black asterisks (*) indicate the channels showing significant contribution to the decoding above chance (p<0.05).

The decoding data thus indicate that the trial-by-trial variability in the neural correlates dissociates the two languages early in time for both picture naming and word reading. To confirm this interpretation, we conducted two further analyses to assess the possibility that this dissociation could be driven by the small but consistent differences in lexical variables (i.e., lexical frequency and length) between the Basque and Spanish stimulus sets, rather than by the different task languages.

First, we performed the same temporal decoding analyses reported above separately for high and low frequency words in each task. Given the lexical characteristics of the high frequency words were highly similar across both languages but showed greater differences for low frequency words (see Table 1), we reasoned that this difference between low frequency words could have driven the ability to decode the language condition in both tasks. For high frequency words, the time course of the temporal decoding effects mirrored the analysis performed with the whole set of stimuli (Supplementary Figure 1). On the other hand, the decoding accuracy of the language of use was not improved when analyzing only the low frequency words, with a delay in the peak effects intervals (starting 257 ms for word reading and 347 ms for picture naming). Despite the between language differences in lexical parameters (log frequency, number of letters and numbers of syllables) of low frequency words, these differences did not improve the decoding accuracy of the language of use. If anything, decoding was earlier and more accurate for higher frequency words, especially in picture naming.

Second, to directly test the potential influence of the lexical parameters on our main findings, we evaluated whether high and low frequency items could be temporally decoded in either language. Importantly, no reliable decoding of high vs. low frequency words was observed in either language for either task (Supplementary Figure 2).

Although there was a smaller number of stimuli (mean trials across participants 39.64 per class) and consequent reduced statistical power in the temporal decoding analyses, these two additional analyses (Supplementary Figures 1 and 2) support the interpretation that language of use is the main factor explaining our results. In addition, it further confirms that lexical differences between the stimuli in each language were not contributing towards the language decoding effects reported in Figure 3a at all.

### Searchlight across sensor-space

Searchlight analysis (Figure 3b) successfully identified the group of sensors contributing to decoding each language in both (covert) word reading and picture naming. For picture naming, the sensors contributing most to the language classification were more lateralized to the right hemisphere, with a dominant cluster in right occipital and temporal sensors. For word reading, the sensors contributing most to language decoding were left-lateralized and the effect was dominant in occipital-temporal sensors, with a smaller cluster in left fronto-temporal sensors. We thus observed a lateralization effect for word reading and picture naming in opposite hemispheres.

### Temporal generalization

Temporal generalization matrices for word reading and picture naming show strong diagonal decoding as reflected by a long-lasting significant cluster. Similar to the time-resolved decoding, the picture naming data showed early reliable language decoding (the cluster appears on the diagonal starting at 87 ms) with a peak decoding AUC value of 0.56 at 392 ms, reflecting a slowly increasing dissociation between the languages across time. Here the generalization plot indicates that the language classifier can most reliably generalize across time points after ∼300-400ms ms.

In the word reading data, the decoding starts a little bit later (117 ms) than in picture naming, but peaks much earlier at 162 ms post-stimulus onset with a peak decoding AUC value of 0.56. After this initial activity a second accuracy peak become evident at ∼400 ms with decoding AUC values around 0.55. In word reading, the classifiers trained at ∼150-160 ms (close to the peak decoding effect) could generalize to the neural activity arising later, ∼250-700 ms after the stimulus presentation.

Temporal generalization thus confirms the pattern of results that emerged in the temporal decoding analyses: a long-lasting language effect for picture naming and the two-stage processing steps for word reading. Generalization plots for word reading indicate that the classifier trained in the early time interval can successfully classify neural data in a later time interval at around 400 ms.^1^

### Cross-task decoding

In the previous section we observed that the neural activity associated to either Spanish or Basque for word reading and picture naming follow slightly different generalization patterns. Thus, we computed cross decoding across picture naming and word reading to estimate if there is any overlap in the neural dissociation between the two languages elicited by the two tasks.

In Figure 5 (left), we report how the classifier trained on picture naming generalizes to the word reading task. An early short-lasting effect is visible, where the picture naming classifier trained at around 77-90 ms generalizes to the neural activity starting from 250 ms post-stimulus onset. A more robust effect emerges later, with the picture naming data from the 200-700 ms time interval eliciting reliable classification for the word reading data in the ∼250-500 ms time interval with an accuracy peak at ∼300 ms.

**Figure 4:**
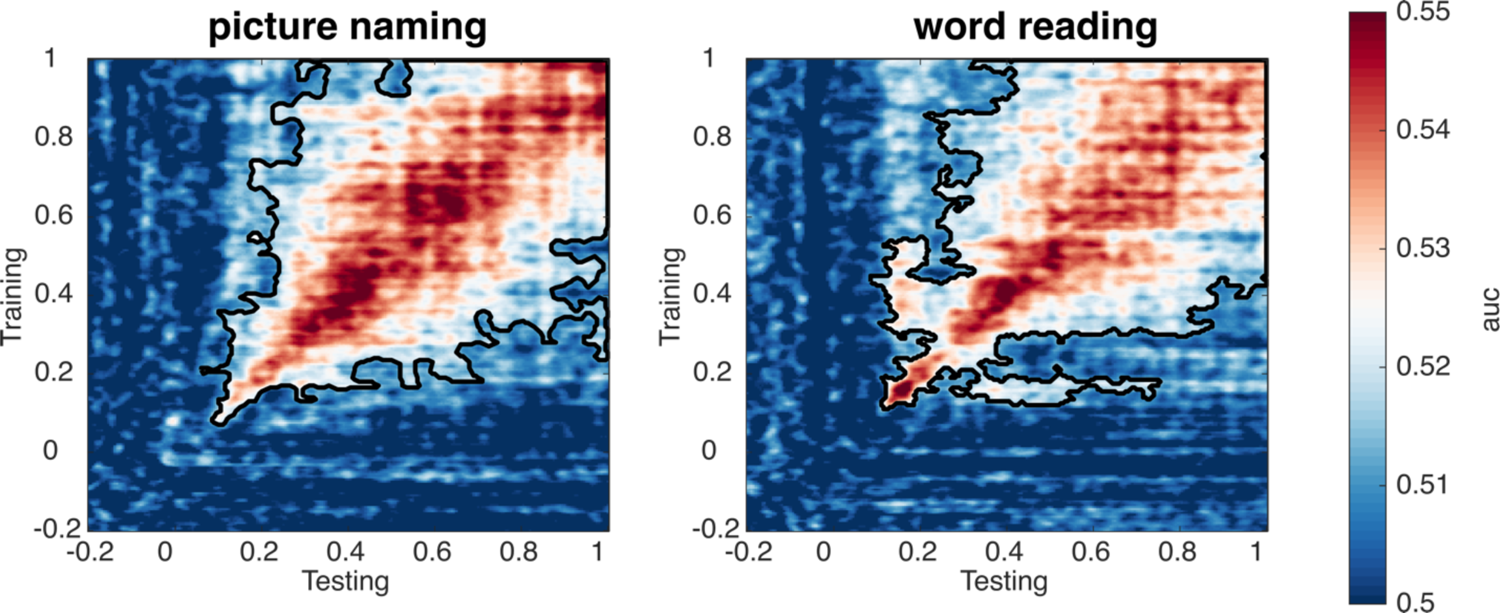
Temporal generalization matrices for language decoding (Spanish vs. Basque, x-axis reports time in seconds) in picture naming (left) and word reading (right). The black lines in each figures indicate the boundary of significant time points emerging from cluster-based permutation tests (p < 0.05).

**Figure 5:**
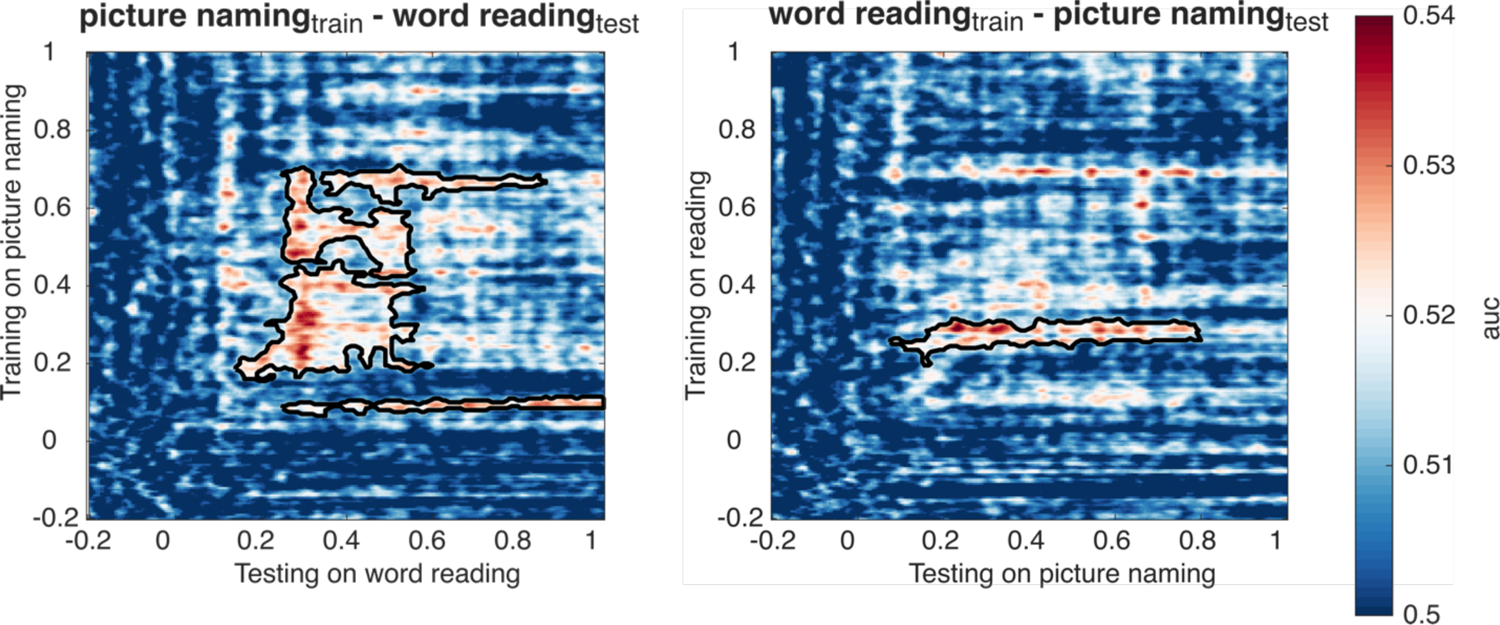
Cross decoding results for language (Spanish vs. Basque, x-axis reports time in seconds). The figure on the left was trained on picture naming and tested on the word reading data, whereas the figure on the right was trained on word reading data and tested on picture naming. The dotted black lines in the figures indicate the boundary of significant time points emerging from cluster-based permutation tests (p < 0.05).

In Figure 5 (right), we report how the classifier trained on word reading generalizes to the picture naming task. Here it is possible to appreciate that the reading data around 230-300 ms triggered reliable language classification in the picture naming task in a long-lasting ∼100-800 ms time interval.

These cross-decoding results confirm that similar language-specific representations are activated in the two tasks, but with different time courses: between 200 and 800 ms in picture naming, and around 300 ms in word reading.

## Discussion

The present study tested the potential of non-invasive neuroimaging for decoding which of two task languages is being covertly produced by bilingual speakers. While invasive brain imaging has highlighted that distinct brain regions could causally modulate production of each language in bilingual speakers (Lucas et al., 2004; Ojemann and Whitaker, 1978), noninvasive brain imaging has been elusive in providing experimental evidence for such dissociation (see Comstock and Oliver, 2021). Overall, we show that (i) language decoding emerges early after stimulus onset (∼100 ms) for both picture naming and word reading; (ii) the effect is sustained in time reaching its peak at ∼430 ms for picture naming and ∼160 ms for word reading; (iii) processing in right occipital-temporal regions is dissociable for the two languages in picture naming, while left occipital regions contribute most to language classification in word reading; (iv) similar language-specific representations are recruited in the two tasks with different temporal dynamics: 200-800 ms for picture naming and around 300 ms for word reading. This pattern of results is not supporting the “integrated” view of the bilingual mental lexicon. On the opposite, bilingual lexical processing seems to recruit dissociable mental representations for the two languages.

One important aspect of the present study was the use of two complementary linguistic tasks, picture naming and word reading. Interestingly, we observed very early language dissociation (∼100 ms) in both tasks when covert production was triggered. This is much earlier compared to classical language-related ERP components, which have been reported starting around ∼200 ms (for single word reading: Kutas and Federmeier, 2011; Swaab et al., 2012; for picture naming: Ganushchak and Schiller, 2009; Indefrey and Levelt, 2004). In the word reading task, one could argue that an early perceptual distinction of the word stimuli in the two languages could have triggered such an early effect (although Casaponsa and Duñabeitia, 2016, reported orthographic effects later in time, ∼250 ms). Importantly, in the picture naming task we used the same images for both languages, meaning that such an early effect cannot be ascribed to visual dissimilarities in the stimulus materials. On the other hand, in picture naming, one could argue that different motor programs are recruited for the two tasks, thus eliciting the early effect. We view this explanation to be unlikely since pre-articulatory effects are typically observed later in time after stimulus onset (> 400 ms: Indefrey and Levelt, 2004). In addition, when the number of syllables was balanced between languages, the language effect was still present and reliable (Supplementary Figure 1). Finally, the phonetic repertoire of Spanish and Basque is almost identical, thus posing similar constraints on the articulatory system.

Overall, the early neural dissociation observed has relevant implications for theoretical accounts of the neural dynamics of language processing. Our data suggest that the divergence of language-specific processing in the bilingual brain occurs hierarchically earlier than the timing associated with linguistic processes, as revealed by previous ERP literature. However, it should be noted that the decoding of conceptual features for both words and pictures (such as, for instance, decoding the animal/tool dissociation, (Dirani and Pylkkänen, 2023; Giari et al., 2020; Leonardelli et al., 2019) has been reported as early as ∼100 ms. The findings that conceptual information already affects neural activity at this early time window supports the possibility that language-specific lexical-semantic representations in the bilingual brain could be available as early as visual decoding processing is completed in preparation for production.

From a linguistic point of view, determining which level of linguistic representation contributed more to the decoding of the language of use is not straightforward based on the present experimental design. In fact, the timings and the generalization patterns (Figures 3 and 4) were not very dissimilar in the two tasks. One difference that could be observed is that in picture naming the decoding accuracy grew slowly, reaching its peak late (> 500 ms), while in word reading the accuracy reached its peak very early (∼170 ms). In previous research, we already showed this asymmetry between the processing of words and pictures in a language task (Giannelli and Molinaro, 2018). Word production models (e.g., Indefrey and Levelt, 2004) assume that pictures first activate lexical-semantic representations that cascade information to sub-lexical (phonological and articulatory) processing later. On the other hand, reading models assume early sub-lexical (visual and orthographic) processing supporting lexical-semantic activation (e.g., Carreiras et al., 2014). Based on this asymmetry, both lexical/semantic and phonological/orthographic representations contribute to successful language decoding, perhaps with slightly better decoding accuracy triggered by sub-lexical features. However, this last claim cannot be fully supported based on the present results since we did not observe any statistical difference in the accuracy level in the two tasks. Nonetheless, we think it is a hypothesis that requires further scrutiny.

A clearer dissociation between the two tasks was observed in the searchlight analyses. While in the word reading task, the posterior and left-lateralized sensors contributed more to language classification, in picture naming the pattern was almost the opposite, with more contribution from right-lateralized sensors. First of all, it is interesting to observe such a strong posterior distribution of such effects, similar to previous MEG studies on picture naming (Geng et al., 2022; Salmelin et al., 1994). Second, while the left-lateralized effect reported for word reading could be predicted based on reading models (e.g., Rueckl et al., 2015), the right-lateralized effect observed for picture naming is less expected. In reading, the involvement of the right hemisphere appears to be mainly related to proficiency levels (Cargnelutti et al., 2019). Since we focused on a group of balanced bilinguals, proficiency should not affect our decoding analysis. On the other hand, activation of different neural representations involved in reading (Dehaene et al., 2015) could be the critical factor boosting decoding accuracy. For picture naming, a critical involvement of the right hemisphere has been discussed in previous fMRI metanalyses (see Comstock and Oliver, 2021; Sulpizio et al., 2020). It is possible that, even if picture naming recruits both hemispheres (Geng et al., 2022), the language-based neural dissociation is stronger in the right-lateralized network. In previous studies, the right temporal regions have been associated with language control (Luk et al., 2011), while the right supramarginal grey matter density was modulated by the vocabulary size of bilinguals (Grogan et al., 2012). Importantly though, the present speculations on why word reading and picture naming recruit different hemispheres during language selection in bilinguals are mainly based on functional and structural MRI data. Evidence from time-resolved decoding of neurophysiological data is scarce. Future research should determine if the task-related hemispheric dissociation we report here reflects a functional reorganization of the two hemispheres based on bilingual experience.

Another clear asymmetry between the two tasks emerged in the cross-decoding analysis, with a more extended effect emerging from the generalization of the picture naming to the word reading task (for a similar pattern of cross-decoding findings, see Dirani and Pylkkänen, 2023). The generalization pattern indicates that brain activity elicited by picture naming between 200 and 700 ms could generalize to brain activity emerging in word reading at ∼300 ms. Interestingly, the opposite was detected in the word reading to picture naming cross-decoding: word reading related brain activity at ∼300 ms generalized to brain activity elicited by picture naming in the 200-800 ms time window. This indicates that language-specific representations that are commonly recruited by both tasks are active for a longer period in picture naming compared to word reading. This cross-task effect can be explained by the different time course in the recruitment of lexical-semantic information. On the one hand, in picture naming, such representations are recruited early and maintained for a long time for the selection of the correct sub-lexical units, even triggering a continuous self-monitoring loop that plays a critical role in language production (Levelt et al., 1999). On the other hand, in word reading, such lexical-semantic representations are activated for a shorter amount of time during the classical N400 time interval. Critically, lexical-semantic information is not critical for reading aloud, since “surface reading” can be successfully achieved without access to the meaning of words (as implemented in the dual route cascaded (DRC) model by Coltheart et al., (2001). This explanation fits with the idea that common language-specific representations in the two tasks are lexical-semantic in nature and are differently recruited depending on the complexity of the language production processes required.

Overall, our study confirms that the two hemispheres are differentially recruited for word reading and picture naming. Importantly, we provide novel evidence for the possibility of detecting language-specific neural correlates in the bilingual mind within these task-related activity patterns. By focusing on a population of Spanish-Basque balanced bilinguals and exploiting the item-specific variability in the neural correlates of two different language tasks, we show that neural decoding analyses can contribute to our knowledge on language-specific processing in the bilingual brain. This bridges the gap between non-invasive and invasive experimental evidence on bilingual lexical processing.

## Acknowledgments

This research was supported by the Basque Government through the BERC 2018–2021 program and by the Spanish State Research Agency through BCBL’s Severo Ochoa excellence accreditation CEX2020-001010-S and the project BES-2016-077560 funded by the Spanish Ministry of Economy and Competitiveness (MINECO). SN acknowledges the support from “The Adaptive Mind,” funded by the Excellence Program of the Hessian Ministry of Higher Education, Science, Research and Art. MC was supported by “la Caixa” Foundation (ID 100010434), under the agreement HR18-00178-DYSTHAL, and by the Agencia Estatal de Investigación PID2021-122918OB-I00. NM was supported by the Spanish Ministry of Science, Innovation and University (grants PSI2015-65694-P, RTI2018-096311-B-I00), the Agencia Estatal de Investigación (AEI), the Fondo Europeo de Desarrollo Regional (FEDER) and by the Basque government (grant PI_2016_1_0014). SN also thank Prof. Daniel Kaiser, Mathematics Institute, JLU Giessen for providing valuable suggestions related to methods used in this manuscript. SN does thank the BCBL lab research staff for their valuable support.

## Author contributions

**NM**: Conceptualization, Methodology, Validation, Supervision, Funding acquisition, Project administration, Writing – review & editing. **SN**: Conceptualization, Methodology, Software, Data curation, Writing – original draft. **MC**: Conceptualization, Methodology, Supervision, Writing – review & editing.

## Data availability statement

The preprocessed dataset is available upon requests directed to Dr. Nicola Molinaro or Dr. Sanjeev Nara. The data set could then be shared through the private BCBL-secured institutional servers temporarily available for big data transfer.

## Supplementary Section

**Supplementary Figure 1:**
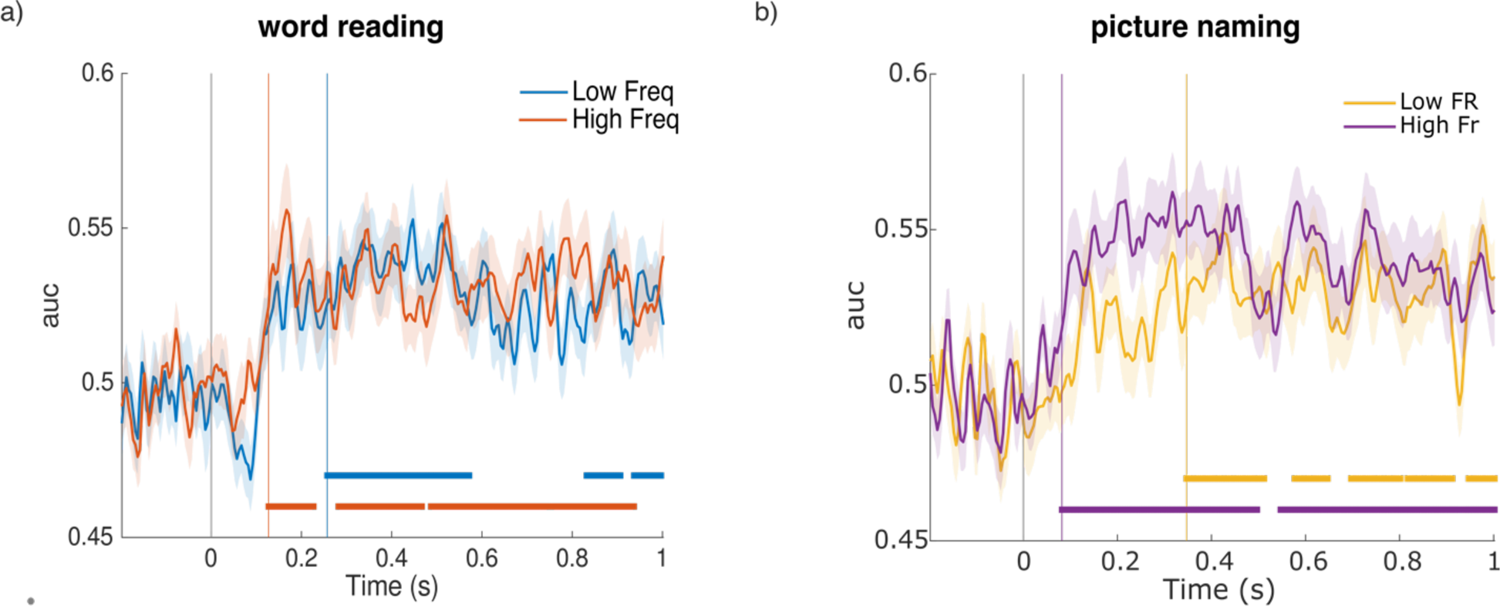
a) Time-resolved language decoding (Spanish vs. Basque) in word reading based on low frequency (blue) and high frequency words (red) separately. The horizontal lines in the figure indicate time points showing significant decoding above chance level (p < 0.05). b) Time-resolved language decoding (Spanish vs. Basque) in picture naming based on low frequency (yellow) and high frequency words (purple) separately. The horizontal lines in the figure indicate time points showing significant decoding above chance level (p < 0.05).

**Supplementary Figure 2:**
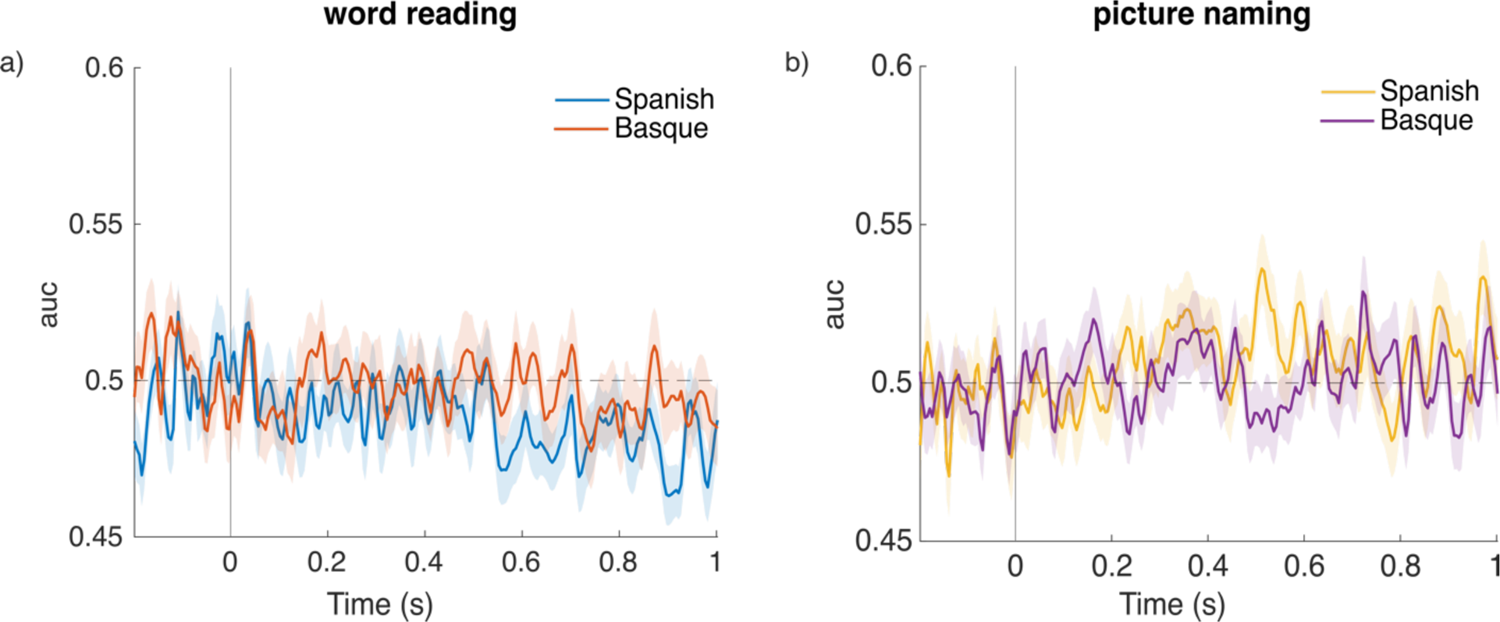
a) Time-resolved language decoding for low vs high frequency words in Spanish and Basque for word reading. b) Time-resolved language decoding the frequency (low vs high freq.) in Spanish and Basque language for picture naming. Note that no time points showed significant decoding above chance level (p < 0.05).

**Supplementary Figure 3:**
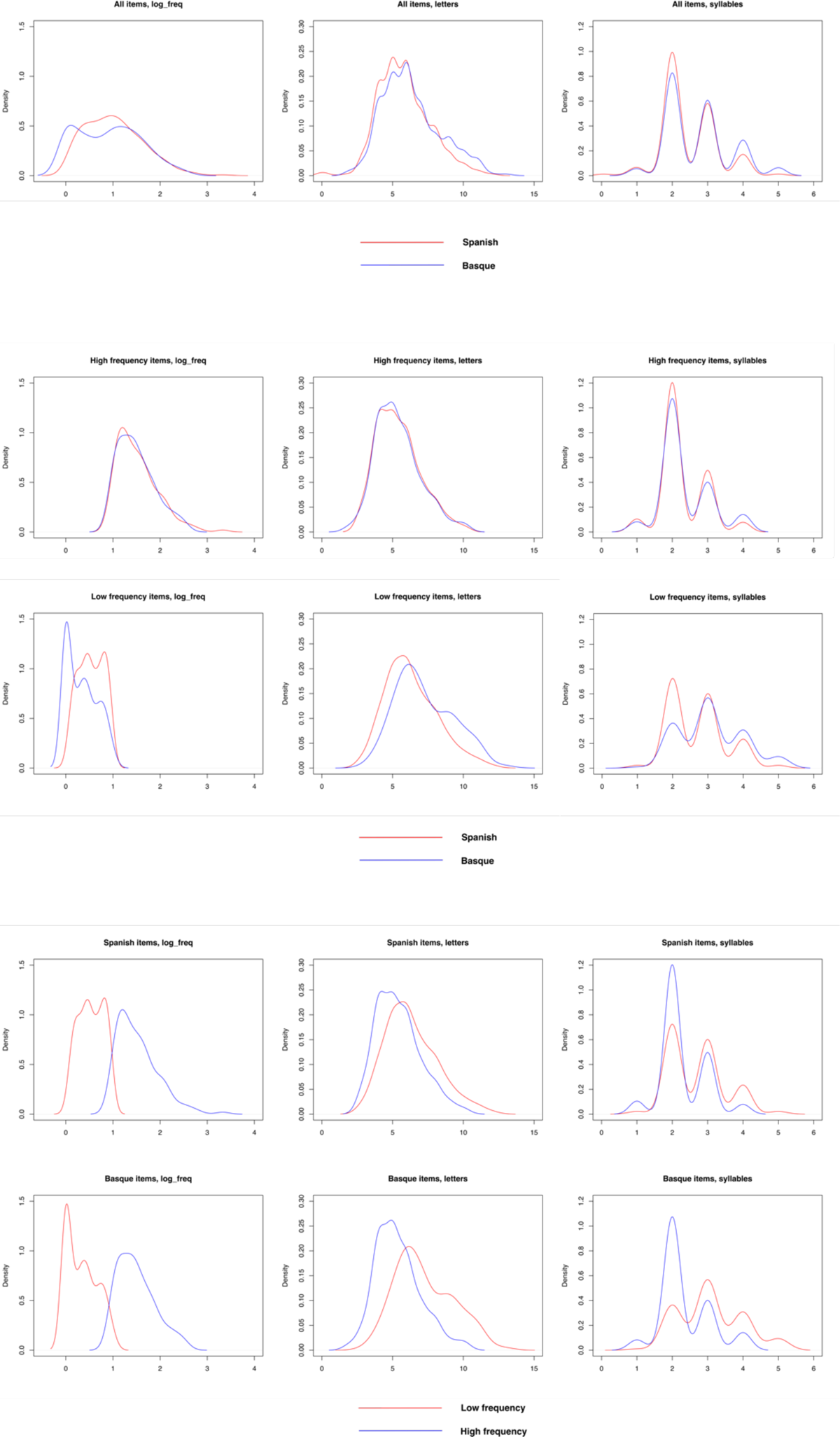
Lexical details of the words used in the experiment. Frequency values based on Spanish (EsPal) and Basque (E-Hitz) language corpora.

## Footnotes

1 We did not perform any additional analysis on potential stimuli subsets, since the number of trials would be too small for the cross-validation procedure. Such additional analyses would be beyond the scope of the generalization analyses, designed to evaluate the organization of the potential brain processes underlying the decoding results (King & Dehaene, 2014).

